# Pollinators, pests and soil properties interactively shape oilseed rape yield

**DOI:** 10.1101/010181

**Authors:** Ignasi Bartomeus, Vesna Gagic, Riccardo Bommarco

## Abstract

1. Pollination, pest control, and soil properties are well known to affect agricultural production. These factors might interactively shape crop yield, but most studies focus on only one of these factors at a time.
2. We used 15 winter oilseed rape (*Brassica napus* L.) fields in Sweden to study how variation among fields in pollinator visitation rates, pollen beetle pest attack rates and soil properties (soil texture, pH and organic carbon) interactively determined crop yield. The fields were embedded in a landscape gradient with contrasting proportions arable and semi-natural land.
3. Pollinator, pest and soil property variables formed bundles across the sites. In general, pollinator visitation and pest levels were negatively correlated and varied independently of soil properties. Because above- and below-ground processes reacted at contrasting spatial scales, it was difficult to predict bundle composition based on the surrounding landscape structure.
4. The above-ground biotic interactions and below-ground abiotic factors interactively affected crop yield. Pollinator visitation was the strongest predictor positively associated with yield. High soil pH also benefited yield, but only at lower pest loads. Surprisingly, high pest loads increased the pollinator benefits for yield.
5. *Synthesis and applications* Implementing management plans at different spatial scales can create synergies among bundles of above- and below-ground ecosystem processes, but both scales are needed given that different processes react to different spatial scales.

## Introduction

Future agriculture needs to be productive to sustain the increasing human population, while conserving biodiversity and the environment. A suggested solution is to stabilize or increase crop yields by maximizing the use of ecosystem services provided by biodiversity, thereby decreasing the dependence on external inputs of agrochemicals in agriculture (Bommarco et al. 2012). However, we don’t fully understand yet how different biotic and abiotic processes interact to shape yield.

Crop pollination is a key ecosystem service that supports crop yield quantity (Garibaldi et al. 2013) and quality (Bartomeus et al. 2014) in three quarters of all crop species (Klein et al. 2007). Another important biotic interaction that determines yield is herbivory by pest insects. They typically reduce yields in all major crops by 5 to 15 percent on average (Oerke and Dehne 2004), and in individual cases yield losses can be far higher (e.g., pollen beetle yield losses in oilseed rape fields may reach up to 80%, Nilsson 1987). Moreover, several soil properties also affect crop production. There is solid evidence from agronomic trials showing that soil texture is associated to water retention (Rawls et al. 1991). Soil organic carbon (SOC) increases the stability of several soil properties (Campbell 1978, Tiessen et al. 1994). Soil pH is closely linked to biological activity in the soil and positively related to nutrient availability and soil fertility (Foth and Ellis 1997), which may translate to higher crop yield (Dick 1992).

Despite the widely acknowledged importance of pollination, pest herbivory and soil properties for shaping yield, the information we have on the joint effects of these factors on yields is fragmentary at best, because they are generally studied in isolation. Hence, processes above- and below-ground are most often implicitly considered as additive in their contribution to crop yield (Bennett et al. 2009). An important practical implication from this is that the management and monitoring of each respective process is considered to be stacked in the landscape. That above- and below-ground processes additively affect plant growth has been challenged in small-scale experiments (Van der Putten et al. 2001, Bezemer et al. 2005). However, at larger spatial scales their interactions remain unstudied (but see Barber et al. 2012) despite above- and below-ground communities can be powerful mutual drivers, with both positive and negative feedbacks (Wardle et al. 2004, Strauss and Irwin 2004).

Pollination has most often been studied as a context-independent process, but recent studies suggest that pollination success and subsequent crop yield are linked to other factors, either via common drivers or through direct interactions between these factors in the yield formation process (Bos et al. 2007, Wielgoss et al. 2013, Classen et al. 2014, Motzke et al. 2014). For example, Lundin et al. (2013) experimentally show that pollinators and pest control of a seed predator interact synergistically, and produce higher yield in combination than the sum of the parts. Local crop management can also interact synergistically with pollination. There is recent evidence that irrigation positively affects the net benefit that plants can take from pollinators in two contrasting crops, coffee and almond (Boreaux et al. 2013, Klein et al. 2014). More generally, it is expected that below-ground soil properties, as well as related ecosystem services provided by soil organisms (Wagg et al. 2014), enhance water retention and nutrient assimilation, and hence should interact with biotic interactions such as pollination and pest damage above-ground (e.g. Williams et al. 2014).

Most evidence about interactive effects on yield between above- and below- ground processes comes from experimental studies. We lack detailed data on how crop yield is affected by multiple processes in agricultural field and at the scales at which crop cultivation takes place – in the arable field and in the surrounding landscape (but see Boreaux et al. 2013). For example, pollinators and natural enemies to crop pests are both affected by landscape composition at scales up to several kilometeres (Shackelford et al. 2013), whereas soil properties are mostly affected locally by management of the individual arable field. Hence, policy-relevant assessments of ecosystem services in agricultural landscapes cannot rely on the simple assumption that a certain land-use results in a given service supply, because not only local field management, but also the composition of the surrounding landscape is an important determinant of biodiversity and ecosystem services (Gabriel et al. 2010). Attempts to maximize the production of a single ecosystem service can result in substantial declines in the provision of other ecosystem services (Bennett et al. 2009, Raudsepp-Hearne et al. 2010).

Here, we use fifteen winter oilseed rape (*Brassica napus* L.) fields situated in a landscape gradient with contrasting proportions of arable and semi-natural land to study natural levels of variation in pollinator visitation rates, pest attack rates and soil properties. We assess the relative importance of each factor for yield formation in an important field crop, as well as potential interactions occurring among them.

## Material and Methods

### Study sites

Fifteen conventional winter oilseed rape (*B. napus,* varieties Excalibur and Compass) fields were selected in 2013 in the Västergötland region, Sweden, along a landscape gradient with contrasting proportions arable and semi-natural land. All sites where located at least 3 km apart from each other. Västergötland is dominated by arable land, mainly cereals, and woodlands, with a small fraction of pastures and meadows. Percentage of arable land was used as a proxy of agricultural intensification (Steffan-Dewenter et al. 2002, Thies et al. 2003, Fahrig 2013) and was measured on multiple scales (see below) using information on land-use characteristics available from the Integrated Administration and Control System (IACS), a data base developed by the Swedish Board of Agriculture. The landscape gradient ranged from 20 to 80 % of arable land in all radii considered. In each field we sampled a non-sprayed area of 40*70 m, situated 30 meters from the edge into the field to avoid edge effects.

### Sampling

Pollinators were sampled twice during peak bloom. For each site and round, we established three 0.5 m^2^ quadrats randomly placed along a 50 m transect centered in the non-sprayed area, parallel to its length. We observed each quadrat for 5 minutes and recorded all pollinators. To record a flower visitor as a pollinator, the insect had to have contact with the central parts of the flower, i.e., the anthers or stigma. Insects were assigned to one of the following categories by visual inspection: Honey bee (*Apis mellifera* L.), bumble bees (*Bombus sp.*), wild bees (diverse species, mostly in the genus *Andrena*), hoverflies (Syrphidae) and other species (mostly Diptera, Hymenoptera and Lepidoptera). All observations were done by a single observer. Pollinators were only sampled on days with sun or scattered clouds and at wind speeds <15 km/h.

Pollen beetles (*Meligethes aeneus* F.), a major pest on oilseed rape (Alford et al., 2003), were counted at four sampling plots 5m apart. Adult pollen beetles were counted on ten plants at each sampling plot (i.e., on 40 plants per field in total). Counts were done three times in the season between the pollen beetle colonization in green bud stage and until flowering was over.

To measure soil properties, we collected five random 15 cm deep soil cores (6 cm diameter) at each site. Cores were mixed and transported at 5°C and protected from sunlight. We determined pH (SS-ISO 10390), proportion of soil organic carbon (SOC) after dry combustion (SS-ISO 10694) and soil texture, measured by determination of percent clay and percent sand particles in mineral soil material after sieving and sedimentation (SS-ISO 11277). All soil analyses were done by Agrilab, Uppsala (http://www.agrilab.se).

Yield was measured as total seed weight per plant just before harvesting. Number of pods was counted on 5 plants per plot, using the same four plots as used for pollen beetles counts (i.e., 20 plants per field). Number of seeds per pod was counted on 20 pods randomly chosen from five plants at each sampling plot (80 pods per field). Weight of 100 seeds from randomly selected pods was measured three times per sampling plot. Yield was measured as total seed weight per plant. It was calculated at the plot level as pods per plant * mean seeds per pod * mean seed weight. We estimated total crop yield as weight of seed obtained per plant, because it integrates fruit and seed set.

### Statistical analysis

First, we identified bundles of above- and below-ground variables potentially affecting yield (analogous to the approach by Raudsepp-Hearne et al. 2010). We ran a K-means cluster analysis on the 15 fields, to identify bundle types, and visualized the results using star plots. Visitation of each pollinator guild, total pest abundance and the three soil properties measured (pH, SOC, and soil texture measured as clay % and sand %) were included in the analysis. We only used one data point per site and variable measured by summing the total number of visits per pollinator guild, or total number of pests across plots and sampling rounds per site. All variables were scaled beforehand to allow meaningful comparisons among variables with different units. The K-means algorithm identifies groupings of observations with similar levels of the included variables. A four-cluster solution was selected to perform the K-means algorithm following a visual assessment of within group sums of squares by number of clusters extracted (Fig. S1 in Supplementary Information). To understand if the clusters of sites with similar levels of above- and below-ground variables are correlated with the landscape structure, we tested if cluster identity is explained by the percentage of agricultural land in the surrounding landscape. We present results for an intermediate scale with a 1500m landscape buffer, but results where qualitatively equal at any radius ranging from 250m to 3km.

Furthermore, we explored at which landscape scale each variable individually responded to the percentage of arable land. Each variable was regressed against percentage agricultural land at increasing radius ranging from 250m to 3km. The most explanatory radius was selected based on maximized r^2^ values.

In addition, we present in the supplementary information pairwise Pearson correlations among all factors measured (Text S1, Table S1) and a principal component analysis (PCA; Fig. S2) which defines an orthogonal coordinate system that optimally describes the variance in our data and that was used to visually represent synergies and trade-offs among the variables.

Second, we assessed the influence of the above- and below-ground factors on crop yield. We used general mixed effects models with crop yield per plant as the response variable and total pollinator visits, pest levels and soil properties as predictors. For each soil property investigated, we used one estimate per field. We pooled all pollinator visits per site; pollinators move freely among plants, and the total visitation abundance in a field is a relevant measure to relate to yield. To avoid over-parametrization of the statistical models, we pooled all guilds and analyzed total visitation because it is a good proxy of pollination (Vazquez et al. 2005, Garibaldi et al. 2013). We used pollen beetle counts per plot because pollen beetles are less mobile and can be patchily distributed (Williams and Ferguson 2010). Finally, we measured the yield from five plants in each plot. Hence, in all models, “plot” nested with in “field” were included as random factor. The full model included the total pollinator visits, the pest counts per plot, and the three soil properties (pH, SOC and clay percent as a measure of texture). We included all pairwise interactions and selected the best models based on AICc (Burnham and Anderson 2002) using the *dredge* function in package MuMin (Barton 2013). We averaged among models within 2 AICc points. All variables were centered beforehand to enhance interpretability of the interactions (Cleasby and Nakagawa 2011). All models were visually inspected for normality of errors and heteroscedasticity. We checked for collinearity in the models by estimating the variance inflation factors (VIF). All VIFs were below 3, hence, there was no strong collinearity in the models. All analyses were done in R, using the base package and nlme (Pinheiro 2014).

## Results

The solution with four clusters was selected as it maximized the variance explained (Fig. S1). However, the other solutions provided qualitatively similar results. The first cluster contained four sites, and was characterized by having lots of hoverflies and high percent of SOC and pH. The second cluster comprised four sites characterized by moderate levels of pests and honeybees, and also wild bees and clay soils. The third cluster was formed by only one site with very high levels of the pest and low pollinator levels. Last, the fourth cluster was comprised by 6 sites, with abundant honey bees and bumble bees, and also dominated by clay soils (Fig. 1).

**Fig 1.**
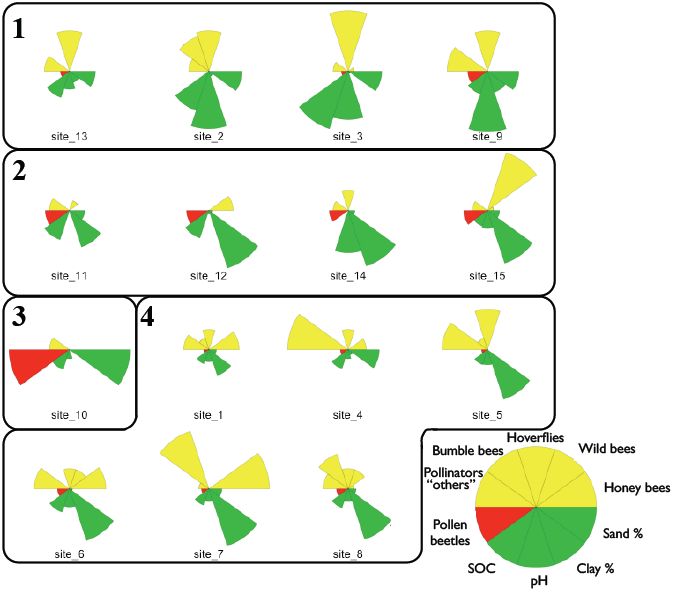
Star diagrams of all 15 sites, showing the 4 clusters of above- and below-ground process identified by the K-means analysis.

Clusters were not explained by landscape structure at any scale (for 1500m radius: F_3,11_ = 1.3, p = 0.3), but cluster number two, comprising four sites, was associated with landscapes with a large percentage of agriculture, while the other two clusters with multiple fields were spread along the agricultural % gradient (Fig. 2). To further explore this disconnection between the bundles observed and the landscape structure, we investigated at which scale each variable responded. As expected, pollinators in general responded negatively to percent of agriculture in the landscape (estimate of total pollinator visits at 3000 m radius = −0.07 ± 0.03, p = 0.03), but guilds responded at contrasting scales; with wild bees responding at very small radius (250 m), while bumblebees and honeybees responded at radii up to 2.5 - 3 km (Fig. 3a). Overall, total pollinator visits response peaked at 3 km radius because honeybees and bumblebees are more abundant than the wild bees. Pollen beetles responded positively to percent agriculture at a scale of 2.5 km (Fig. 3b), but the trend is not significant (estimate = 4.1 ± 2.29, p = 0.09). None of the soil properties was significantly affected by the percentage of arable land at any scale (Fig. 3c; all models p > 0.2).

**Fig 2.**
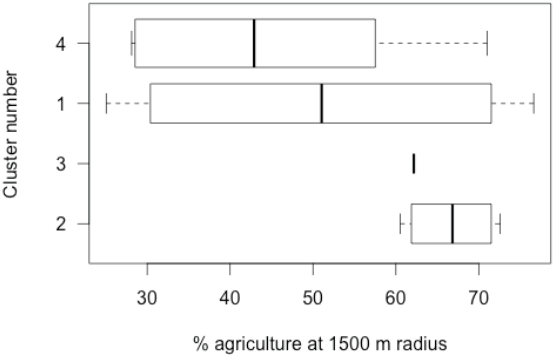
Relationship between the 4 bundles identified by the cluster analysis and the percentage of agriculture in the landscape. Although cluster 2 is associated with more agricultural areas, there is no overall pattern relating those bundles to the underlying landscape structure.

**Fig 3.**
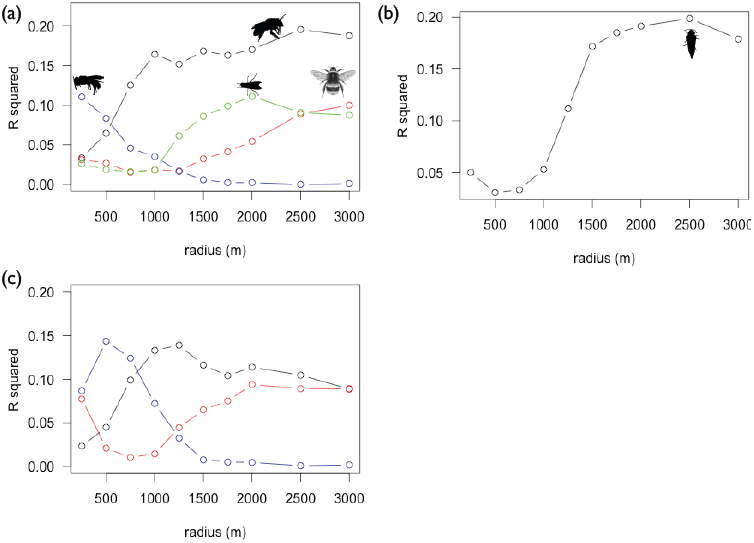
Explanatory power of percent of agriculture in the landscape at different scales for A) Pollinators (honey bee in black, wild bees in blue, bumble bees in red and hoverflies in green), B) Pollen beetles and C) Soil properties (Total organic carbon in Black, pH in red and % clay in blue).

The PCA reflected the clustering pattern and showed that overall, sites with lower pest levels tended to have more pollinators, and that those variables are independent of soil properties (Fig. S2).

When analyzing the effect on yield, we found seven models within two AICc points (Table S2) with pollinators, pH and pests retained in most models. The averaged model (Table 1) shows that pollinators are positively correlated with yield and that there is an interaction with the pest, such that at high pest numbers, the relationship with pollinators is steeper (Fig. 5a). This interaction should be interpreted with care, given that there are few data points with high levels of both, because they are weakly, but negatively correlated (VIF < 3). Interestingly, pH only had a positive effect on yield when pest levels were low, but at high pest levels, the relationship disappears (Fig. 5b). The best model marginal r^2^ is 0.20, while the conditional r^2^ is 0.55 (Nakagawa 2013).

**Table 1:**
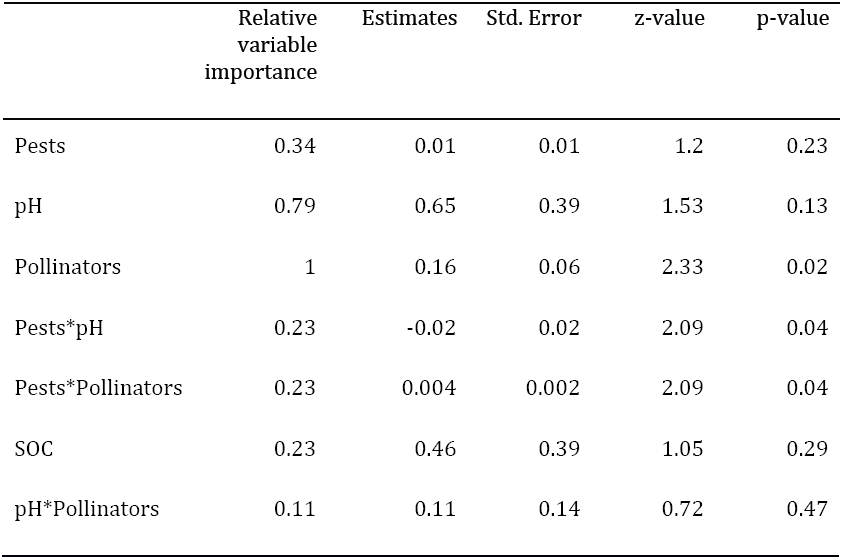
Model-averaged coefficients of the model predicting oilseed rape yield. The relative importance indicates the proportion of models containing each predictor, being “Pollinators” the only variables retained in all models.

| Relative variable importance | Estimates | Std. Error | z-value | p-value |
| --- | --- | --- | --- | --- |

|  |  |  |  |  |  |
| --- | --- | --- | --- | --- | --- |
| Pests | 0.34 | 0.01 | 0.01 | 1.2 | 0.23 |
| pH | 0.79 | 0.65 | 0.39 | 1.53 | 0.13 |
| Pollinators | 1 | 0.16 | 0.06 | 2.33 | 0.02 |
| Pests*pH | 0.23 | -0.02 | 0.02 | 2.09 | 0.04 |
| Pests*Pollinators | 0.23 | 0.004 | 0.002 | 2.09 | 0.04 |
| SOC | 0.23 | 0.46 | 0.39 | 1.05 | 0.29 |
| pH*Pollinators | 0.11 | 0.11 | 0.14 | 0.72 | 0.47 |

**Fig 4.**
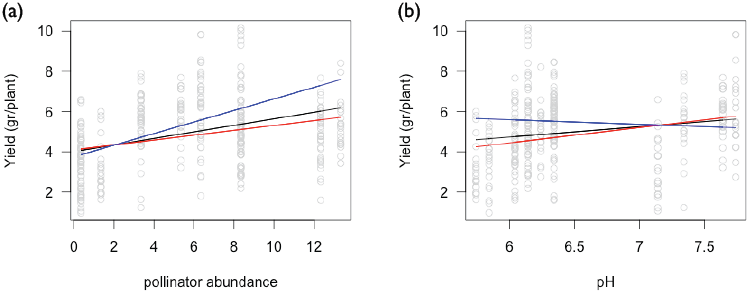
Relationship of A) pollinators and b) pH with yield. Black lines are estimate predictions for the average level of pests. Red lines are predictions for low and blue lines for high levels of pests respectively.

## Discussion

Crop yield is shaped by combinations of biotic and abiotic factors. Identifying the main above- and below-ground factors for assuring high yield requires an examination of how they naturally co-vary in the landscape, as well as a simultaneous estimation of several potential drivers. We show that pollination, pest levels and soil properties (mainly soil pH) are key factors for winter oilseed rape yield formation. Although these have been independently identified as important for yield formation in a number of crops (Garibaldi et al. 2013, Oerke et al. 2006, Dick 1992), their individual correlations with yield are usually low. For instance, even if there is a robust general trend of increasing yield with increasing pollinator visitation there is a great deal of unexplained variation (Garibaldi et al. 2013), and sites with similar pollinator levels often differ substantially in yield. Studies addressing several ecosystem services and abiotic factors simultaneously have the potential to explain more of this variation. Importantly, we show that such factors can interact, thereby modifying the outcome of the main effects. Hence, our study adds to recent experimental evidence that the response of yield to one factor or resource such as pollination depend on other variables such as pest control levels, and that their effects are not additively contributing to yield (Lundin et al. 2013). However, in our dataset, even after accounting for pollinator visits, pest attack rates and several soil properties, the fixed factors predicts only a 20 % of the variance, while the random factors associated with unmeasured field variables explain up to 55%.

We identified four bundle types among our explanatory variables, indicating that certain variables tend to occur together (e.g., honey bees, bumble bees and clay soils in cluster 4). However, these are not predicted from the landscape characteristics in which the target fields were embedded. More generally, pollen beetles and the most abundant pollinators (i.e., honey bees and bumble bees) naturally co-varied negatively with each other. This negative correlation between pollen beetles and pollinators is partially explained by the landscape analysis, as both respond to percent of arable land at similar large scales, but in opposite directions. One explanation for this pattern is that pollinators respond positively to an increased amount of feeding and nesting resources in complex landscapes (Kennedy et al. 2013), and that pollen beetle abundances are lowered by natural enemies that also are benefited by such landscapes (Chaplin-Kramer et al. 2011). However, given that pollen beetles feed on flower buds and are still active on flowers during the pollination period, they can also have a direct effect by deterring pollinators from heavily infested fields. Interestingly, we show an interaction between pollen beetles and pollinators. Contrary to expected, at the same pollinator visits level, the pollinators’ positive effect on yield is higher when abundances of pollen beetles are high. Hence, rather than pollen beetles lowering the visitation efficiency (e.g., by reducing pollen availability) or directly damage the plant (e.g., increasing fruit abortion rates; Alford et al. 2003), it seems that the observed pollen beetle damage to buds may result in considerable compensatory growth by oilseed rape. For example, it has been reported that moderate feeding damage to the terminal raceme leads to increased production of new side racemes (Williams and Free 1979, Tatchell 1983, Lerin 1987, Axelsen and Nielsen 1990). It is interesting that this compensatory growth is only beneficial under high pollination, and may indicate that the benefit may only arise if this newly produced branches are well pollinated.

We also show that soil properties vary across sites, independently to the proportion arable land in the landscape. Soil pH seems to be the most important soil factor explaining yield in our analyses. Interestingly, the positive effect of soil pH on yield is only detectable at low pest levels. This implies that at high pest levels, the benefits from increasing pH and thereby soil fertility are not translated into increased yield, but may instead be lost to pest damage or invested into plant defenses. In fact, soil fertility can increase plant defenses (Coley et al. 1985) and we found that fields with a high pH tended to have rather low pest levels. This pattern was weak, but was found both in the cluster analysis and in the PCA (Fig. S2).

Surprisingly, soil texture (i.e., proportion clay), which is positively related to water retention and nutrient exchange capacity, was not retained in any of the best models explaining yield. This indicates that water was probably not a limiting factor in this year and region. However, clay contents variable may be important in years with low precipitation, and for other climatic regions or crops (see Boreaux et al. 2013, Klein et al. 2014).

As expected, soil properties were not affected by the percent of arable land in the surrounding landscape (Williams et al. 2013), and hence they co-vary independently with pollination and pests. This implies that management practices to sustain yield are needed both at the field as well as in the wider surrounding landscape. Few studies have simultaneously considered effects of local (on field) and landscape scale land use on multiple ecosystem functions (Bianchi et al. 2006).

Our results support recent claims that interactions among ecosystem services are to be expected, but the importance of the key above- and below-ground variables affecting yield and their interactive effects are likely to be crop specific and to vary between sites and years. For example, the degree of plant dependency on pollinators will determine the potential benefit that can be achieved by pollinators. However, even in plants with high rates of self-pollination, yield quality is enhanced with insect pollination (Bartomeus et al. 2014). Herbivores that affect the reproductive parts of the plant, such as seed weevils (Lundin et al. 2013) or pollen beetles (this study) are more likely to directly interact with the benefits from pollination. Herbivore plant suckers or defoliators can be nutrient sinks that affect fruit formation, even when sufficient pollination is achieved (Bos et al. 2007). Plant species-specific pathways to absorb, assimilate and mobilize nutrients will determine how above- and below-ground factors interact. For example, coffee plantations can trigger one or two flowering peaks a year clearly affecting pollinator responses, and this depends on nutrient and water availability (Boreaux et al. 2013). More studies on a variety of cropping systems and ecosystems including abiotic and biotic variables are needed in order to reach any generality.

The strength and shape of the relationships between different above- and below-ground processes is poorly known. This is partly because we lack information about synergies and trade-offs in the management of multiple processes. We show that interactions between biotic and abiotic factors can give rise to scale-dependent synergies when managing multiple ecosystem services. Hence, both above-ground biotic interactions regulated at large scales and below-ground abiotic factors managed at local scales interact to form crop yield.

Data analyzed: uploaded as online supporting information

## Acknowledgments

Field work was conducted by Oskar R. Rubbmark, Gerard Malsher and Laura Riggi. Audrey St-Martin helped with the soil samples. Ola Lundin provided valuable comments on an earlier draft of this paper. Funding was provided by the Swedish research council FORMAS to R.B.

## Supplementary Information

### Text S1: Correlation among variables

Pearson correlations among the pairwise variables studied are usually low with some exceptions. Among the pollinators, honey bees and bumble bees were positively correlated (r = 0.47, p = 0.07). Similarly, some belowground properties are correlated. As expected, sand and clay percent are negatively correlated (r = -0.85, p < 0.001) and SOC is negatively correlated with clay percent (r = - 0.54, p = 0.04). Moreover, hoverflies are correlated with several soil properties (SOC r = 0.50, p = 0.06; pH r = 0.69, p = 0.004; Clay = -0.47, p = 0.08) and with pest levels (r = 0.54, p = 0.04). Finally, pests are correlated with sand percent (r = 0.48, p = 0.06).

The first two axes of the PCA explained together 55% of the variance (31% and 24% respectively; Fig. S2), with subsequent axes explaining less than 15% each. We found a trade-off between pests and pollinators, with sites with lower pest levels (loadings on second axes = -0.76), having more pollinators (loadings in second axes honeybees = 0.63 and bumblebees = 0.62). The less abundant wild bees and hoverflies are independent of honeybee and bumblebee visits, and co-vary in opposite directions among them (loadings in first axes = -0.49 and 0.93, respectively). This uncoupled responses among pollinators is the base for a possible biodiversity insurance against environmental fluctuations. Along the first axes, total organic carbon and pH correlate well (loadings on first axes = 0.61 and 0.72 respectively) and partially sand content (loading in first axes = 0.34, but also -0.79 in second axes). As expected, clay content follows an opposite trend as sand content (loading in first axes = -0.64, but 0.63 in second axes).

**Fig. S1.**
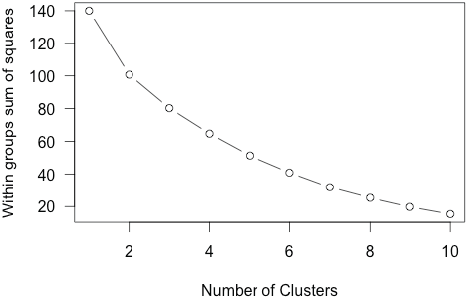
Scree plot showing the within groups sum of squares as a function of the number of clusters selected.

**Fig. S2.**
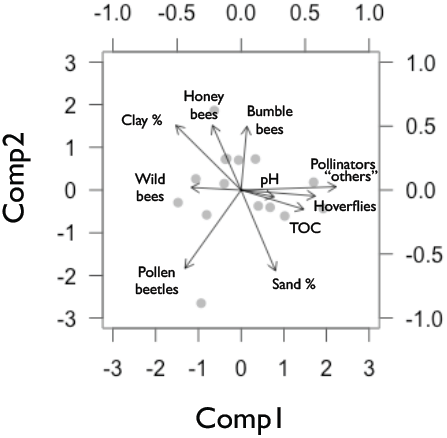
First two axes of the principal component analysis. PCA loadings: Honey bee (PC1 = -0.28, PC2 = 0.63), Wild bees (PC1 = -0.49, PC2 = 0.03), Hoverflies (PC1 = 0.93, PC2 = 0.03), Bumble bees (PC1 = 0.06, PC2 = 0.62), Other pollinators (PC1 = 0.32, PC2 = -0.07), Pollen beetles (PC1 = -0.55, PC2 = -0.76), SOC (PC1 = 0.61, PC2 = -0.19), pH (PC1 = 0.72, PC2 = -0.06), Clay percent (PC1 = -0.64, PC2 = 0.63), Sand percent (PC1 = 0.34, PC2 = -0.79).

**Table S1.** Full correlation table, upper triangle has the p-values, lower triangle the Pearson r correlation values.

|  | A.mellifera | Wild bees | Syrphids | Bombus | Other | M.aeneus |
| --- | --- | --- | --- | --- | --- | --- |
| A.mellifera | 1.00 | 0.88 | 0.16 | 0.07 | 0.59 | 0.34 |
| Wild bees | -0.04 | 1.00 | 0.19 | 0.72 | 0.66 | 0.65 |
| Syrphids | -0.37 | -0.35 | 1.00 | 0.75 | 0.20 | 0.03 |
| Bombus | 0.47 | -0.10 | -0.08 | 1.00 | 0.47 | 0.15 |
| Other | -0.15 | -0.12 | 0.34 | -0.20 | 1.00 | 0.41 |
| M.aeneus | -0.26 | 0.12 | -0.54 | -0.38 | -0.22 | 1.00 |
| SOC | -0.14 | -0.15 | 0.49 | -0.04 | -0.16 | -0.19 |
| pH | -0.34 | -0.17 | 0.69 | 0.01 | 0.03 | -0.26 |
| Clay percent | 0.38 | 0.39 | -0.47 | 0.21 | -0.29 | -0.09 |
| Sand percent | -0.35 | -0.26 | 0.15 | -0.13 | 0.09 | 0.48 |
|  | SOC | pH | Clay percent | Sand percent |  |  |
| A.mellifera | 0.60 | 0.20 | 0.16 | 0.20 |  |  |
| Wild bees | 0.58 | 0.54 | 0.14 | 0.34 |  |  |
| Syrphids | 0.05 | 0.00 | 0.07 | 0.57 |  |  |
| Bombus | 0.86 | 0.96 | 0.43 | 0.61 |  |  |
| Other | 0.55 | 0.90 | 0.27 | 0.73 |  |  |
| M.aeneus | 0.48 | 0.33 | 0.73 | 0.06 |  |  |
| SOC | 1.00 | 0.24 | 0.03 | 0.16 |  |  |
| pH | 0.32 | 1.00 | 0.22 | 0.31 |  |  |
| Clay percent | -0.53 | -0.33 | 1.00 | 0.01 |  |  |
| Sand percent | 0.38 | 0.28 | -0.85 | 1.00 |  |  |

**Table S2.** Complete list of models within 2 AICc points

|  | (Int) | pest | pH | pollinators | SOC | pest:pH | pest:pol | pH:pol | df | logLik | AICc | delta | weight |
| --- | --- | --- | --- | --- | --- | --- | --- | --- | --- | --- | --- | --- | --- |
| 1551 | 5.12 | 0.013 | 0.48 | 0.22 |  | -0.022 | 0.004 |  | 9 | -543.199 | 1105.0 | 0.00 | 0.230 |
| 13 | 4.95 |  | 0.69 | 0.14 |  |  |  |  | 6 | -546.411 | 1105.1 | 0.09 | 0.219 |
| 29 | 4.95 |  | 0.59 | 0.15 | 0.37 |  |  |  | 7 | -545.928 | 1106.2 | 1.23 | 0.125 |
| 15 | 4.95 | 0.004 | 0.77 | 0.16 |  |  |  |  | 7 | -546.063 | 1106.5 | 1.49 | 0.109 |
| 9 | 4.93 |  |  | 0.13 |  |  |  |  | 5 | -548.163 | 1106.5 | 1.52 | 0.108 |
| 4109 | 4.99 |  | 0.87 | 0.17 |  |  |  | 0.11 | 7 | -546.086 | 1106.5 | 1.54 | 0.107 |
| 25 | 4.94 |  |  | 0.14 | 0.56 |  |  |  | 6 | -547.172 | 1106.6 | 1.62 | 0.102 |

